# The *Drosophila* TNF Eiger contributes to Myc super-competition independent of JNK activity

**DOI:** 10.1101/2021.04.06.438672

**Authors:** Albana L. Kodra, Aditi Sharma Singh, Claire de la Cova, Marcello Ziosi, Laura A. Johnston

## Abstract

Numerous factors have been implicated in the cell-cell interactions that lead to elimination of cells via cell competition, a context-dependent process of cell selection in somatic tissues that is based on comparisons of cellular fitness. Here we use a series of genetic tests in *Drosophila* to explore the relative contribution of the pleiotropic cytokine Tumor Necrosis Factor ⍺ (TNF⍺) in Myc-mediated cell competition (also known as Myc super-competition or Myc cell competition). We find that the sole *Drosophila* TNF, Eiger (Egr), its receptor Grindelwald (Grnd/TNFR), and the adaptor proteins Traf4 and Traf6 are required to eliminate wild-type “loser” cells during Myc cell competition. Although typically the interaction between Egr and Grnd leads to cell death by activating the Jun N-terminal Kinase (JNK) stress signaling pathway, our experiments reveal that many components of canonical JNK signaling are dispensable for cell death in Myc cell competition, including the JNKKK Tak1, the JNKK Hemipterous (Hep) and the JNK Basket (BSK). Our results suggest that Egr/Grnd signaling participates in Myc cell competition, but functions in a role that is independent of JNK activation.

## Introduction

Cell competition is an evolutionarily conserved process of somatic cell selection that has been implicated in numerous biological processes, including embryonic development, tissue maturation, organ size control and cancer (CLAVERÍA *et al*. 2013; SANCHO *et al*. 2013; ELLIS *et al*. 2019; SUN *et al*. 2023; TSHERING *et al*. 2023; ZHANG *et al*. 2023). Cell competition leads to the elimination via death or expulsion of viable, but relatively weak cells (“losers”) from growing tissues, allowing more robust cells (“winners”) to populate the tissue (JOHNSTON 2009). The specific mechanisms that recognize fitness differences and trigger elimination of the loser cells are influenced by the organism, tissue type, genetic background, and the particular circumstances of the competitive environment. First identified in *Drosophila,* cell competition employs context-dependent activation of signaling pathways involved in regulation of stress, immunity or apoptosis. Signaling by the Jun N-terminal kinase (JNK) stress response pathway is often stimulated (MORENO *et al*. 2002a; DE LA COVA *et al*. 2004; MORENO AND BASLER 2004; TYLER *et al*. 2007; IGAKI *et al*. 2009; OHSAWA *et al*. 2011; KUCINSKI *et al*. 2017; BANRETI AND MEIER). In some competitive contexts in *Drosophila*, components from the innate immune response are activated, forming a novel signaling module consisting of select components of the Toll and the Immune Deficiency (IMD) signaling pathways (hereafter called the **CCSM**, for **Cell Competition Signaling Module**) (Supp. Fig. 1). The CCSM mediates the competitive interactions between wildtype (WT) cells and cells expressing an extra copy of the conserved transcriptional regulator Myc, leading to the former’s death and elimination from the tissue (MEYER *et al*.; ALPAR *et al*. 2018; NAGATA *et al*.; HOF-MICHEL *et al*.). In this work we focus on interactions between the CCSM and signaling from the TNF/TNFR pathway.

The CCSM and the IMD and Toll immune response pathways share several molecular components, but there are notable differences. In contrast to the canonical IMD and Toll pathways, the CCSM requires only a subset of factors from each pathway. Moreover, the CCSM is not activated in immune tissues such as the fat body (FB), and does not activate a typical immune gene expression program. In Myc cell competition, genetic requirements have been demonstrated for several Toll related receptors (TRRs), the secreted Toll ligand Spätzle (Spz), and 2 proteases necessary for the processing of the pro-Spz protein (MEYER *et al*.; ALPAR *et al*.), but the highly conserved TRR adaptor Myd88 is dispensable (MEYER *et al*. 2014). From the IMD pathway, the adaptor FADD, the caspase-8 homolog Dredd, and the NF-kB factor, Relish (Rel) are genetically required, but IMD itself is not, nor is the Rel-phosphorylating IKK complex. Notably, expression of Dredd is sufficient to induce Rel cleavage in S2 cells (STOVEN *et al*. 2000; STOVEN *et al*. 2003; ERTURK-HASDEMIR *et al*. 2009; PAQUETTE *et al*. 2010), and *in vivo*, Dredd expression induces Rel-dependent cell death (HU AND YANG; CHINCHORE *et al*. 2012; KIM *et al*. 2014; MEYER *et al*. 2014). In addition, the CCSM is activated locally in Myc-mediated cell competition, within the mosaic tissue, in response to the confrontation of WT and Myc-expressing cells and its activity remains restricted to the competing cells (ALPAR *et al*.). Activation of the CCSM occurs as a result of transcriptional up-regulation of the Spz pro-protein and the serine protease Spz-processing enzyme (SPE) in the Myc super-competitor cells. Production of these proteins leads to the proteolytic cleavage of Spz into an active ligand, allowing it to associate with one or more of 5 different TRRs that appear to be highly expressed in the wild type (WT) cells. This appears to restrict activation of the CCSM to the WT cells, converting them into “losers” that induce Rel/NF-kB-dependent expression of the Hid proapoptotic factor, and ultimately to their death (MEYER *et al*. 2014; ALPAR *et al*. 2018). The CCSM thus consists of a novel configuration of immune response components that activate a genetic program distinct from that of the conventional *Drosophila* immune response (MEYER *et al*. 2014; ALPAR *et al*. 2018).

Reporters of the JNK stress pathway such as *puc-lacZ*, an enhancer trap insertion in the *puckered* (*puc*) locus (encoding a phosphatase antagonist of JNK activity) (MARTIN-BLANCO *et al*. 1998; MCEWEN AND PEIFER 2005) indicate that JNK activity is present in some epithelial cells of the wing and eye imaginal discs during their normal growth, where it promotes cell survival by blocking JNK activity (MCEWEN AND PEIFER 2005; WILLSEY *et al*.). It has also been observed in wing discs during Myc super-competition, where *puc-lacZ* can be variably activated in either loser or winner cells (DE LA COVA *et al*.; MORENO AND BASLER). However, the importance of JNK activity in loser cell elimination in Myc cell competition has remained unclear. In one report, only 30% of loser cell death was suppressed by the absence of JNK signaling (DE LA COVA *et al*.). In another, suppression of loser cell death by blocking JNK signaling also required expression of p35, a pan-caspase inhibitor (MORENO AND BASLER). By contrast, the death of WT loser cells in Myc super-competition is completely suppressed by the genetic loss of any individual component of the CCSM (Supp. Fig. 1) (MEYER *et al*.).

JNK signaling that leads to cell death is induced by activation of the highly conserved and pleiotropic Tumor Necrosis Factor α (TNFα) signaling pathway in mammals and in *Drosophila* (NAGATA 1997; IGAKI AND MIURA 2014; COLOMBANI AND ANDERSEN 2023). The sole *Drosophila* TNF, known as Eiger (Egr), has homology with mammalian Ectodysplasia (EDA)-A2 and other TNF ligand superfamily proteins (e.g. RANKL, CD40L, FasL, TNF-α, and TRAIL) (IGAKI *et al*. 2002). Like other TNFs, Egr is a type II transmembrane protein that also can be cleaved and glycosylated, allowing it to be secreted (KAUPPILA *et al*. 2003). Egr binds to the TNF receptors (TNFRs) Wengen (Wgn) (KANDA *et al*.; KAUPPILA *et al*.) and Grindelwald (Grnd) (ANDERSEN *et al*. 2015). The Egr/TNFR complex, via the Traf 4 and Traf 6 adaptors, activates the core kinase cascade of the JNK pathway (Supp. Fig. 1), which includes the MAP3K/JNKK kinase Tak1, the Tak1-associated binding protein 2 (Tab2), the MAP2K/JNK kinase Hemipterous (Hep), and the MAPK/JNK Basket (Bsk) (Supp. Fig.1)(KANDA *et al*.; KAUPPILA *et al*. 2003; ANDERSEN *et al*. 2015). Egr/TNFR signaling is activated by numerous kinds of stress, including the innate immune response, where it cooperates with the IMD pathway to activate Rel-mediated expression of anti-microbial peptides (AMPs) in the FB, an adipose tissue and major immune organ in *Drosophila* larvae (DELANEY *et al*. 2006). Under these conditions, Tak1 is a critical mediator of Rel activation, which requires both proteolytic cleavage by the caspase-8 homolog, Dredd, and phosphorylation by the IkB Kinase (IKK) complex (STOVEN *et al*. 2003; ERTURK-HASDEMIR *et al*. 2009). Tak1 is therefore a key signaling component shared by the IMD/Rel pathway and the core JNK cascade triggered by Egr/TNFR (DELANEY *et al*. 2006; KLEINO AND SILVERMAN). Interactions between JNK signaling and the Toll immune pathway have also been described (WU *et al*. 2015).

With the goal of shedding light on the relative contribution of the JNK signaling pathway in Myc-induced cell competition, in this work we used genetic tests to understand its role in the death and elimination of the WT “loser” cells. Since Egr activates cell death through JNK signaling (ANDERSEN *et al*. 2015), we investigated the role of Egr and its receptors Grnd and Wgn in the activation of JNK signaling during Myc cell competition. We show here that genetic loss of Egr or its receptor Grnd, but not loss of the Wgn/TNFR, robustly blocks the death and elimination of loser cells in competition with cells that express more Myc. Genetic experiments using cell competition assays suggest that Egr/Grnd activity in the WT loser cells requires the function of Traf4 and Traf6. However, our results reveal that downstream of these adaptors, canonical JNK signaling is not necessary for loser cell elimination in this competitive context: genetic experiments indicate that Tak1 and the downstream JNK kinase Hep are dispensable, and a dominant negative form of the kinase Bsk only partially suppresses the competitive elimination of the WT loser cells. Thus although Egr/Grnd function is required in the WT loser cells for their death, their activity is largely independent of JNK signaling. Instead, our results suggest that Egr/Grnd influences the Rel activator Dredd in the killing of WT loser cells. We propose that Egr/Grnd signaling participates in Myc super-competition by promoting the activity of the CCSM.

## Materials and Methods

### Fly strains and Husbandry

Flies were raised at 25°C on standard cornmeal-molasses food (R-food, LabExpress) supplemented with fresh dry yeast. The following strains were used: *hep^r75^, traf4^ex1^*, *wgn^Pe00637^*, *grnd^M105292^*, *grnd^DfBSC149^, Tak1^1^*, *Tak1^2^, UAS-Bsk^DN^*, *rnGal4*, and *act>y+>Gal4* (from Bloomington Drosophila Stock Center, BDSC); *TRE-dsRed 16* (gift of D. Bohmann); *wgn^KO^*, *Egr::GFP* (fTRG) and *Egr-lacZ* (gifts of M. Milan), *egr^3AG^,* (KODRA *et al*.); *UAS-Grnd-IR^KK^* (Vienna Drosophila Resource Center, VDRC); *UAS-egr^IR^* and *UAS-dTRAF4^IR^*(gifts of M. Miura); *UAS-egr^#5^/CyO* and *UAS-dTRAF6^IR^*(gifts of T. Igaki); *UAS-grnd^extra^* and *UAS-grnd^intra^*(gifts of P. Leopold); *UAS-Dredd* (B. Lemaitre); *tub>myc^STOP^>Gal4* (DE LA COVA *et al*.); *tub>CD2^STOP^>Gal4* (MORENO AND BASLER).

### Cell competition assays

Competitive and non-competitive control clone assays were performed in wild-type, homozygous mutant or hemizygous mutant backgrounds as indicated (DE LA COVA *et al*.; DE LA COVA *et al*.; MEYER *et al*.; ALPAR *et al*.). Eggs from appropriate crosses were collected on freshly yeasted grape plates for 2-4 hours (or as indicated in figures) and allowed to develop at 25°C in a humid chamber for 24 hours. After hatching, larvae were transferred to food vials supplemented with fresh yeast paste at densities of less than 40 larvae per vial to prevent crowding. To measure clonal growth and cell competition in a manner that allows precise temporal control of transgene expression, we used FLP recombinase-mediated, intra-molecular recombination (reviewed in (GERMANI *et al*.)). To generate competitive clones, a *tub>myc^STOP^>Gal4* cassette (where > represents a FLP-recognition target (FRT) site) was used to generate random GFP-marked *tub>Gal4* clones (DE LA COVA *et al*. 2004). Prior to recombination, all cells express the *Myc* cDNA under control of the *tubulin* promoter, at a level less than 2-fold over endogenous *myc* levels (WU AND JOHNSTON 2010). Recognition of the FRTs by FLP leads to excision of the *myc* cDNA and transcriptional STOP sequence and generates cells that heritably express Gal4 (*tub>Gal4*) and can regulate expression of UAS-genes of interest (e.g., GFP). As the *GFP-*expressing cells no longer express extra *myc*, they are subject to competition from surrounding cells that retained the *>myc^STOP^>* cassette (DE LA COVA *et al*. 2004). FLP, under heat shock (HS) control, was activated in larvae carrying this cassette by HS at 37°C for 10 minutes, at 48 hours after egg laying (AEL). Post-HS, larvae were allowed to grow at 25°C for 24, 48 or 96 hours, as indicated for each experiment. In parallel, to randomly generate GFP-marked clones as controls for noncompetitive clonal growth, larvae carrying the transgenic cassette, *act>y+ ^STOP^>Gal4* (on chromosome 3) or *tub>CD2^STOP^>Gal4* (on chromosome 2) were used. The control clones were induced with a HS at 37°C for 6 min with the same timing as for the competitive Myc cassette (DE LA COVA *et al*.; DE LA COVA *et al*.; MEYER *et al*.; ALPAR *et al*.). These heat shock times were optimized for each cassette to generate only few clones per disc, to avoid merged clones. A detailed protocol is available upon request.

### Tissue dissection, fixation and Imaging

Larval carcasses containing mosaic wing discs were dissected and inverted and fixed in 4% paraformaldehyde in phosphate-buffered saline (PF-PBS) for 20 min at room temperature and washed with 0.01% Tween-20 in PBS (PBTw). Hoechst 33258 (Sigma) or DAPI (Invitrogen) was used to stain DNA. Wing discs were mounted in VectaShield Antifade on glass slides. Images were acquired with a Zeiss Axiophot with Apotome, Zeiss LSM800, or Leica LSM710 confocal microscope. Clone area (in square pixels or microns) was measured with Image J software. Clones were scored in the central area of the wing disc (wing pouch and proximal hinge), where competition is most severe (ALPAR *et al*.). Nonparametric Mann-Whitney and Kruskal-Wallis tests were used to determine statistical significance as indicated in each figure.

### Immunohistochemistry

Primary antibodies used: guinea pig anti-Grnd (1:200, gift of P. Leopold), rabbit anti-Caspase-3 (1:100, Cell Signaling), rabbit anti-GFP (1:2000, ThermoFisher) and rabbit anti-Dcp-1 (1:100, Cell Signaling) were incubated at 4°C overnight with dissected wing discs previously permeabilized with 0.5% Triton X-100 in PBS (PBTx) for 1 hour and then blocked in PBTw with 5% normal serum for 1 hour at RT. The samples were incubated with the secondary antibodies Alexa555 and Alexa488 (1:600, Molecular Probes) (pre-absorbed against fixed embryos) for 3 hours at RT in the dark and then washed with PBTw either 3x at RT or overnight at 4°C. Hoechst 33258 or DAPI (Sigma) was used to stain DNA. Images were processed and where applicable, assembled from Z-stacks as maximum projections, with Image J software.

### RNA *in situ* hybridization of wing discs

Dissected, fixed and washed larval carcasses were added to 500ul of 300mM ammonium acetate and 500ul of 100% ethanol and gently mixed, washed in 100% ethanol for 10 min at room temperature, washed in xylene/ethanol (1:1) for 10 min, washed in ethanol 3 times, 10 min each, washed in methanol for 2 min, washed in methanol/4% PF (1:1) for 2 min, fixed again in 4% PF-PBTw for 10 min, washed in PBTw 5 times for 5 min each, and incubated at RT in hybridization solution/PBTw (1:1) for 15 min; this was replaced by hybridization solution and incubated at the appropriate hybridization temperature for at least 1hr. Digoxygenin-RNA probes were added to hybridization solution, mixed well, and denatured at 90°C for 5 min followed by incubation on ice for 5 min. The hybridization solution was removed from the carcasses and replaced with the denatured probe in hybridization solution, and incubated at the appropriate temperature overnight. Samples were washed and processed with anti-Digoxygenin antibodies as in Alpar et al (ALPAR *et al*.). A detailed protocol is available upon request.

## Results

### The TNF Egr contributes to the elimination of WT loser cells

Reporters of JNK signaling activity are weakly activated in normally growing WT wing discs (MCEWEN AND PEIFER; HARRIS *et al*.)), and have also been observed in a variety of conditions that lead to cellular stress. Indeed, JNK activity is important in a variety of contexts of cell competition (IGAKI *et al*. 2009; OHSAWA *et al*. 2011; IGAKI AND MIURA 2014). JNK activity has also been observed in Myc super-competition, although in this context, the extent of its contribution to the loser cell fate has remained unclear (DE LA COVA *et al*. 2004; MORENO AND BASLER 2004). The core components of the JNK pathway (Supp. Figure 1) – Traf4, Traf6, Tak1, Tab2 and Hep – are all expressed in wing disc cells (Modencode). Engagement of Egr with either of its receptors, Wgn and Grnd (Figure 2, A-B), allow Egr to be a powerful, JNK-dependent inducer of cell death in the developing eye and wing (IGAKI *et al*. 2002; MORENO *et al*. 2002b). This is illustrated by expression of *egr* in GFP-marked cell clones in wing discs (Fig. 1A-B), which leads to the death and elimination of most of the GFP-positive cells in the WP within 24 hours (Fig.1 B). Many dying cells, visualized using an antibody specific to cleaved and activated Caspase-3 (Cas-3), are visible in a basal focal plane of the wing disc (Fig. 1B’). To assess the role of Egr in Myc cell competition, we examined its endogenous expression in the wing disc using a reporter strain carrying an *egr::GFP* fusion protein (SANCHEZ *et al*. 2019). The growing wing disc epithelium consists of a monolayer of columnar cells (called the disc proper) that is contiguous to and covered by a squamous cell layer called the peripodium (TRIPATHI AND IRVINE 2022). The wing disc also houses the adult myoblast precursor (AMP) cells, which lie beneath the two epithelial layers in the notum (future adult dorsal thorax), and the tracheal primordial cells (TRIPATHI AND IRVINE 2022). During the disc’s rapid growth period (L2-mid-L3), Egr is highly expressed in the non-epithelial AMP and tracheal cells (BROWN *et al*.; CASAS-VILA *et al*.; EVERETTS *et al*.) (Fig. 1E-F), but in the wing pouch (WP) epithelium, which gives rise to the adult wing blade, its expression is seemingly random and limited to a few cells primarily in the “transition zone”, cuboidal cells that lie at interface between the peripodial and columnar layers (MCCLURE AND SCHUBIGER 2005); Fig. 1C, D and data not shown). Egr is also expressed in many other tissues including the FB, from which its soluble form is secreted into the circulating hemolymph and accessible to wing discs and other larval organs (AGRAWAL *et al*. 2016; DE VREEDE *et al*. 2022).

**Figure 1.**
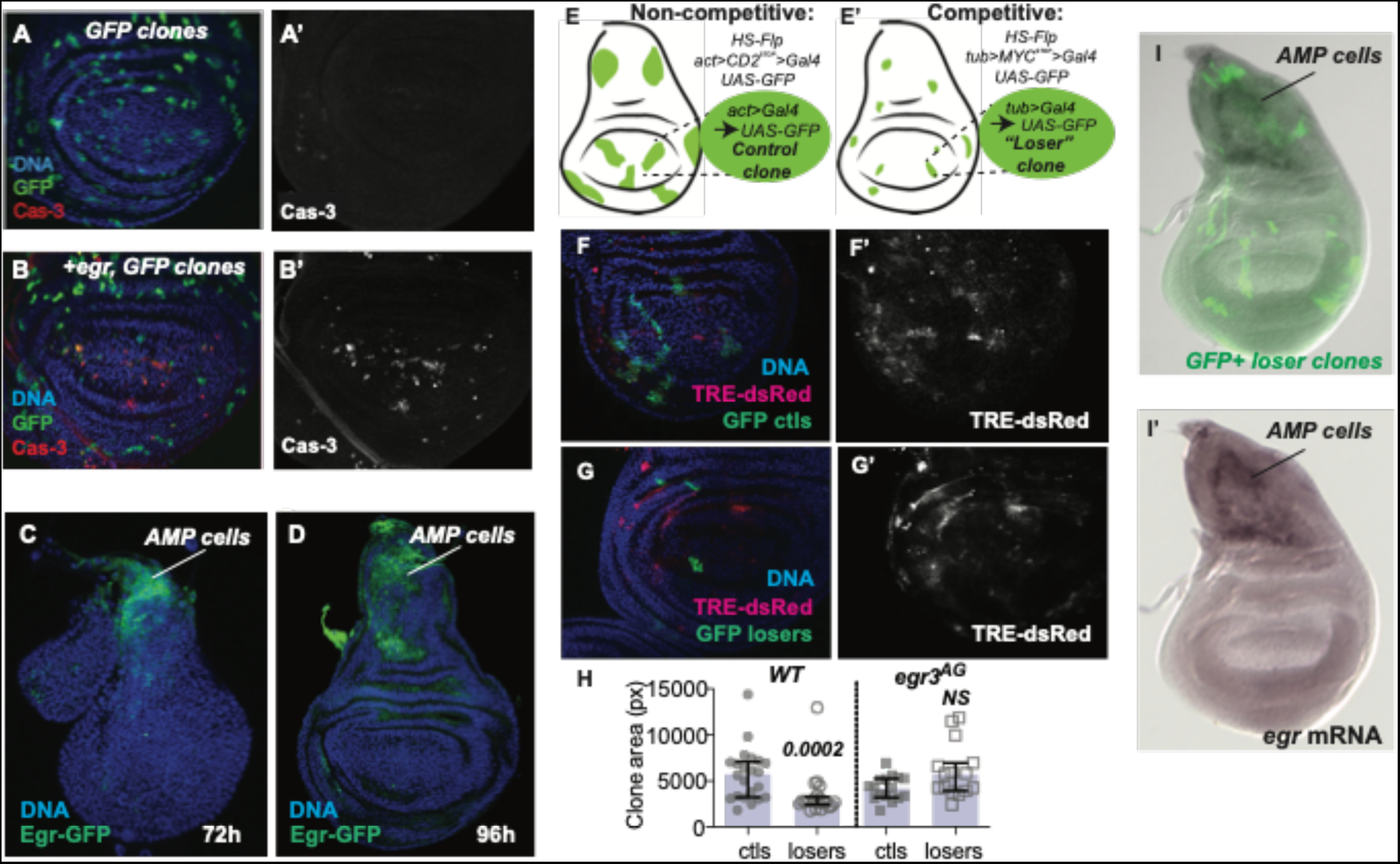
Eiger (Egr) induces cell death and is required for the competitive elimination of WT loser cells. **A-B)**. Control, GFP-expressing clones induced in wing discs and allowed to grow for 24 hrs. **A’)**. Anti-activated Cas-3 immunostaining. **B)**. Clonal expression of *egr* for 24 hrs leads to caspase activity (red) and rapid elimination of clones from wing discs; **B’)**. Anti-activated Cas-3 staining. **C-D)**. Egr::GFP is highly expressed in the AMP cells (arrows) but is very low or undetectable in the rest of the wing disc. **E-F)**. Schematic of Myc cell competition assay. **E)**. Flp-FRT recombination is used to generate non-competitive control clones that express UAS-GFP and allowed to grow for defined periods of time, after which measurement of clone area yields information on clonal growth. **F)**. Similarly, GFP-marked WT clones are generated in a background in which cells express an extra copy of Myc, yielding competitive interactions between the clones and surrounding Myc expressing cells. See Methods for details. **G).** The *egr^3AG^* null allele suppresses the elimination of loser wing disc cells. Clones grew for 50 ± 2 hrs. Error bars show median and interquartile range. Statistical significance was determined by non-parametric Mann-Whitney tests, NS = not significant. **H).** *egr* mRNA *in situ* hybridization in *tub>myc^STOP^>Gal4* wing discs. WT loser clones are marked by GFP expression (H-H’). *egr* mRNA is easily detected in AMP cells in the notum (arrow) but is not detectable in the loser clones, nor in the surrounding *tub>myc^STOP^*expressing cells.

**Figure 2.**
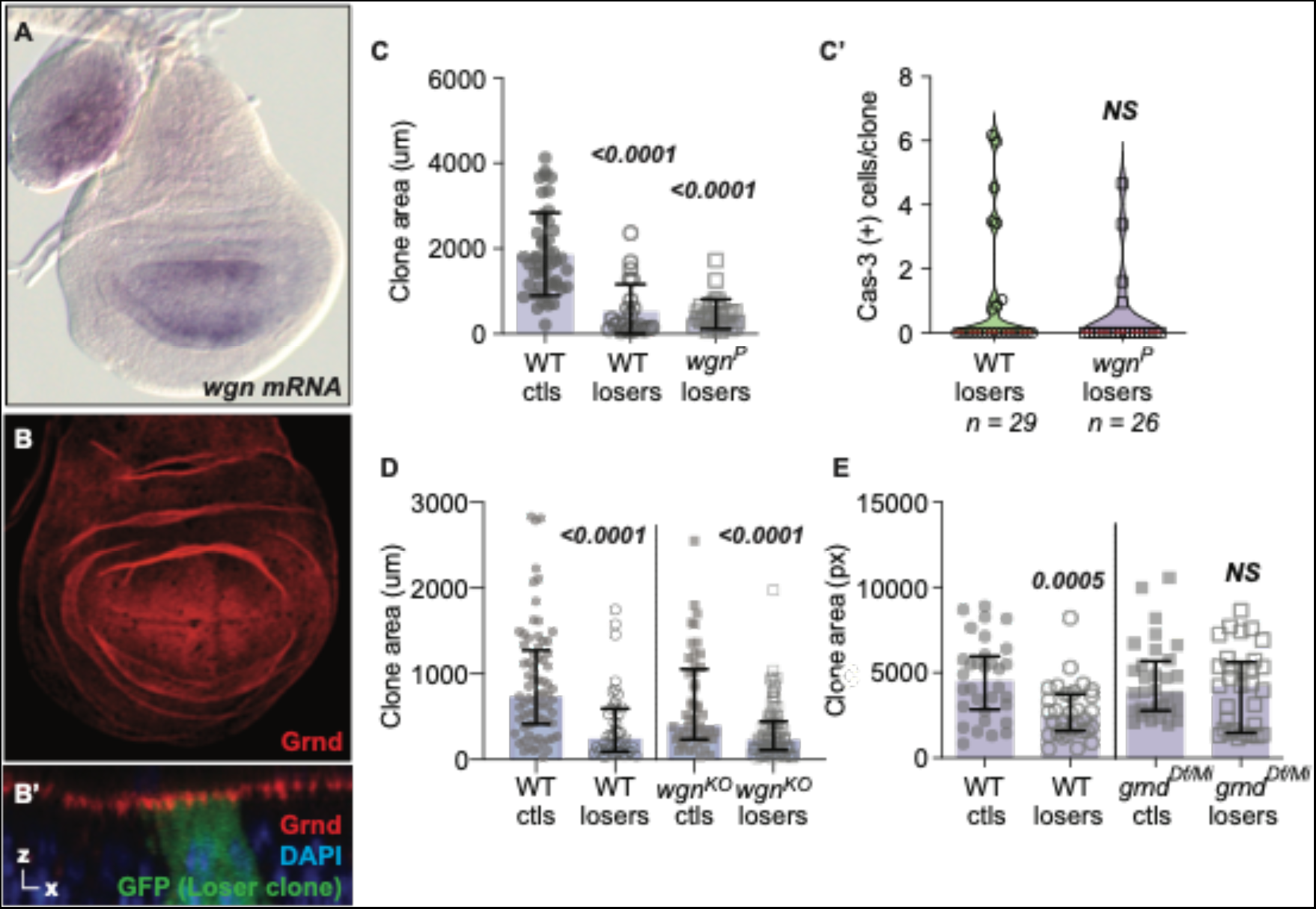
The TNFR Grnd contributes to the elimination of loser cells. **A).** *wgn* mRNA is expressed primarily in the wing pouch (WP) region of the mid-L3 wing disc (90hr AEL). **B).** Grnd protein is uniformly expressed and apically localized in wing disc cells (disc is 120hr AEL; a maximum projection of a z-series is shown). **B’**). Cross-section showing apical Grnd localization in wing disc with GFP-marked WT loser clone (in the *tub>myc^STOP^ >Gal4* background). **C).** *wgn* loss does not suppress loser cell elimination. Loser clones in both WT male and in hemizygous *wgn^P^* male larvae grow to smaller sizes compared to WT control clones. **C’).** Loss of Wgn does not prevent loser cell death. Violin plots of active Cas-3 positive cells measured 24hr after clone induction (ACI) in WT and *wgn^P^* mutants. Wing discs from male larvae were scored. n = number of clones analyzed per genotype. **D).** Competition assay comparing control and loser clones in a WT background with those generated in parallel in *wgn^KO^* mutants. Error bars show median and interquartile range. Clones grew for 48 ± 2 hrs. Cell clones were measured in wing discs from male larvae (*WT/Y* and *wgn^KO^/Y*). **E).** Loss of *grnd* (using the *grnd* null mutants *grnd^Df/^grnd^Mi^)* in *trans* suppresses the elimination of loser cells under competitive conditions, but does not alter control clone size in the non-competitive context. Clones grew for 48 ± 1.5 hrs. Error bars represent median and interquartile range. Statistical significance in **C-E** was determined by non-parametric Mann-Whitney and Kruskal-Wallis tests. NS = not significant.

To investigate the role of Egr in activation of JNK in wing discs during Myc cell competition we examined expression of the JNK-specific *TRE-dsRed* reporter, consisting of multimerized AP-1 binding sites under control of a basal promoter (CHATTERJEE AND BOHMANN 2012) in the context of our Myc cell competition assay (Fig. 1E, F-G). The basis of this cell competition assay is a transgenic FRT cassette, *tub>myc^STOP^>Gal4* (see Methods for details; Fig. 1E-E’) that when intact, drives *myc* expression slightly higher than its endogenous level (<2-fold) (WU AND JOHNSTON). Flp-mediated excision of the >*myc^STOP^*sequences allows heritable expression of Gal4 (*tub>gal4*) and activation of UAS-GFP. Cells in the *tub>gal4* clones are essentially WT for Myc, yet are at a competitive disadvantage due to the non-clonal surrounding cells that continue to express elevated Myc from the intact transgene. This disadvantage leads cells in *tub>Gal4* clones to die at increased frequency (DE LA COVA *et al*. 2004; MORENO AND BASLER; DE LA COVA *et al*. 2014; MEYER *et al*. 2014; MERINO *et al*.), resulting in “loser” clones (Fig. 1E’) that grow to significantly smaller sizes than non-competitive clones generated from a control cassette in parallel (e.g., Fig.1E, WT Ctls). The growth disadvantage and resulting smaller size of the loser clones is due to induction of the proapoptic gene *hid* in loser cells, mediated by the CCSM (DE LA COVA *et al*. 2004; MORENO AND BASLER; DE LA COVA *et al*. 2014; MEYER *et al*. 2014). Using this assay to induce heat-shock (HS)-generated clones, we observed weak, sporadic activation of *TRE-dsRed* in wing discs in both control (non-competitive) and competitive contexts, with no direct correlation of TRE activity in either control or loser clones (Fig. 1F-G). These results indicate that JNK activity is present under these conditions at low levels in the wing disc epithelium, but shows no overt connection to a cell’s specific competitive status.

To assess whether Egr contributed to the death of the loser cells, we tested whether the *egr^3AG^* null allele (KODRA *et al*.) altered the fate of loser cells in the competition assay. In a WT background of cells expressing *tub>myc^STOP^>Gal4*, excision of the cassette led to generation of GFP-marked loser clones that were competitively eliminated from the wing disc, leading to a significant reduction in clone size compared to non-competitive control clones generated in parallel (Fig. 1H, WT ctls and losers). However, in the *egr^3AG^* background, the *egr* mutant clones generated in both competitive and non-competitive contexts grew at the same rate and to a similar final size (Fig. 1H, *egr^3AG^* ctls and losers), indicating that competition was abolished between the *tub>myc^STOP^>* expressing cells and the clones of WT cells. Contrary to expectations, we were unable to detect *egr* transcripts within the loser cells in competition assays by RNA *in situ* hybridization (Fig. 1I-I’) or Egr protein using the Egr::GFP reporter, suggesting that Egr is produced elsewhere in the wing disc or from another tissue in the larva (data not shown; Sharma Singh, in prep). These results indicate that *egr* is genetically required for the elimination of loser cells in wing discs in the *tub>myc^STOP^>Gal4* background, but in this context it does not function in an autocrine manner.

### Grnd/TNFR is required to eliminate WT loser cells from wing discs

The *Drosophila* genome encodes two TNFRs, Wgn and Grnd, both of which are abundantly expressed in wing discs. Egr binds to both Wgn and Grnd, and can activate downstream signaling that requires Traf4 and Traf6 through each receptor (IGAKI *et al*. 2002; KANDA *et al*. 2002; GEUKING *et al*. 2005; ANDERSEN *et al*. 2015). Previous reports of *egr* over-expression in the eye disc linked Egr activity to Wgn, the adaptors Traf4 and Traf6, and cell death via the JNK signaling pathway (IGAKI *et al*. 2002; KANDA *et al*. 2002; GEUKING *et al*. 2005). Although Egr expression in the wing disc is largely restricted to the non-epithelial AMP cells, *wgn* mRNA is robustly expressed in the WP (Fig. 2A), the region of the disc subject to the most severe effects of cell competition (ALPAR *et al*. 2018). As *wgn* is located on the X chromosome, we examined clone growth and cell competition in wing discs from WT and *wgn^P^* mutant male larvae. As expected, after a 48hr growth period, loser clones in WT wing discs were significantly smaller than non-competitive control clones (Fig. 2C, WT ctls vs WT losers), and loser clones in wing discs from *wgn^P^* mutants were outcompeted by *tub>myc^STOP^>Gal4* cells just as efficiently as in WT wing discs (Fig. 2C), with similar numbers of dying cells per clone (Fig. 2C’)). As the *wgnP* allele is likely hypomorphic, we also tested *wgn^KO^*, a targeted knock out allele (ANDERSEN *et al*. 2015). Similarly, loser cells were efficiently eliminated in the competition assay when *wgn* was completely lacking (Fig. 2D). These results suggest that Wgn is not required for eliminatino of WT loser cells, and are consistent with recent work reporting that Wgn has limited function in Egr signaling in wing imaginal discs (PALMERINI *et al*. 2021).

In contrast to Wgn, the TNFR Grnd is highly and uniformly expressed in the wing disc epithelium, localized to sub-apical epithelial cell membranes (Fig. 2B, B’). Although present at high levels, loss of Grnd does not affect cell viability under normal conditions (ANDERSEN *et al*. 2015). When examined in cell competition assays, Grnd protein localized normally to the apical surface of cells in both loser and control clones of third instar larval wing discs (Fig. 2B’). To test whether Grnd functioned in cell competition, we used the trans-heterozygous combination *grnd^Df^/grnd*^Minos^, which expresses no detectable Grnd protein (ANDERSEN *et al*. 2015). Strikingly, the absence of *grnd* suppressed much of the elimination of the loser cells, allowing loser clones to grow to sizes similar to the controls (Fig. 2E, *grnd^Df/Mi^* ctls vs losers). In contrast, *grnd* loss had no effect on the size of non-competitive control clones (Fig. 2E, *grnd^Df/Mi^* ctls vs WT ctls), indicating that Grnd’s role in loser clones was specific to the competitive context. To determine whether Grnd was required autonomously within the loser cells, we expressed *UAS-grnd-RNAi* (*grnd^IR^*) in the loser cells. Again, knock-down of *grnd* impaired cell competition, but had no effect on the growth of clones in a non-competitive environment (see Fig. 5F). Together, our results suggest that Egr functions with Grnd to promote the elimination of the loser cells in wing discs. Because we found no evidence of Egr in the loser or winner cells themselves, we infer that Egr is produced elsewhere in the wing discs or in another larval tissue. We suggest that in this competitive context, Egr functions as a paracrine factor and uses Grnd as its receptor on wing disc cells.

Grnd binds Egr via its extracellular N-terminal CRD domain, which results in signaling transmitted through its intracellular C-terminal domain (ANDERSEN *et al*. 2015). However, expression of only the intracellular portion of Grnd*, UAS-grnd^intra^*, is sufficient to activate JNK signaling and apoptosis in an Egr-independent manner (ANDERSEN *et al*. 2015). Apoptosis induced by expression of *UAS-grnd^intra^* is efficiently suppressed in the *hep^r75^* mutant background, a severe loss of function allele of the Hep/MKK7 protein that is required to transmit JNK signaling (GLISE *et al*.; ANDERSEN *et al*. 2015), confirming that Grnd functions upstream of the JNK signaling cascade. To determine whether Grnd-mediated intracellular signaling is sufficient to eliminate loser cells in cell competition we expressed *UAS-grnd^intra^*in wing discs. As a control, we expressed *UAS-grnd^extra^*, a transgene containing only the extracellular portion of Grnd, which binds to Egr but does not activate JNK signaling (ANDERSEN *et al*. 2015). Expression of *UAS-grnd^extra^*using *rnGal4*, which drives expression in the WP, led to only a slight, patchy rise in cell death (Fig. 3A, A’), consistent with previous results (ANDERSEN *et al*. 2015). *grnd^extra^* also had no effect on growth when expressed in clones, in either a non-competitive or a competitive environment (Fig. 3B, C). In contrast, *rnGal4*-directed expression of *grnd^intra^* led to massive cell death, substantially reducing the size of the wing discs (Fig. 3D, D’). When we expressed *grnd^intra^*in GFP-marked cell clones, despite being initially detectable (Fig. 3E-F insets), wing discs were completely devoid of GFP-positive cells by 48hr ACI, indicating that all of the clones were eliminated (Fig. 3E-F). Thus, our results confirm that expression of *grnd^intra^*, and by inference Grnd activation, transmits a potent death-inducing signal within cells (ANDERSEN *et al*. 2015). Moreover, complete elimination of *grnd^intra^*-expressing clones from the wing epithelium was the outcome in both a non-competitive (Fig. 3E) and a competitive environment (Fig. 3F).

**Figure 3.**
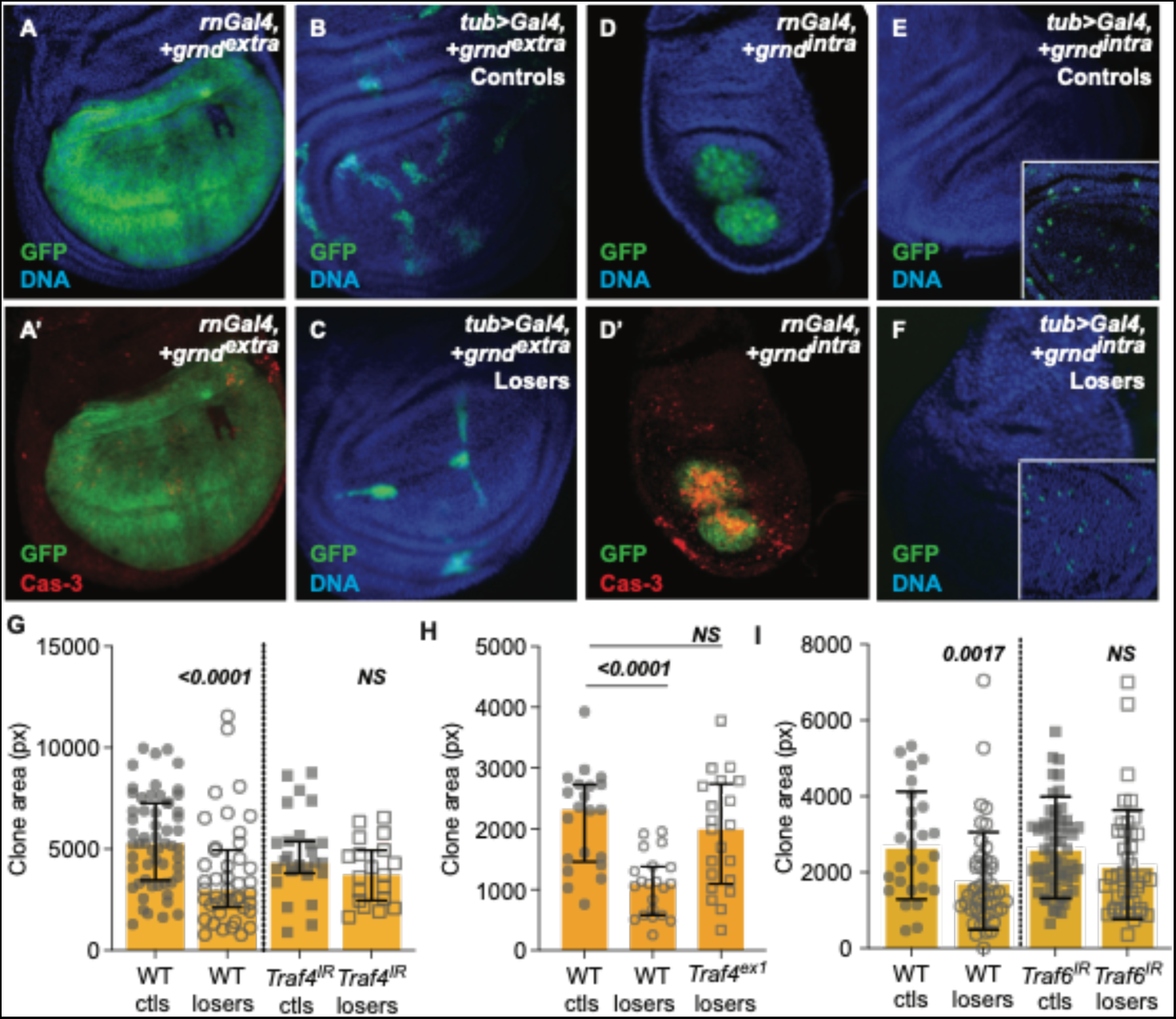
Traf4 and Traf6 contribute to the elimination of loser cells. **A).** Expression of *grnd^extra^* in the wing pouch induces a small number of Cas3-positive cells **(A’)** but otherwise does not alter disc morphology. **B).** Representative wing disc showing non-competitive control clones co-expressing *GFP* and *grnd^extra^*. **C).** Representative wing disc showing loser clones in the competitive context co-expressing *GFP* and *grnd^extra^*. **D).** Expression of *grnd^intra^*induces massive cell death **(D’)** and severely reduces overall wing disc size. **E, F).** Representative wing disc with *grnd^intra^* –expressing clones in non-competitive control **(E)** and competitive backgrounds **(F)** clones 48 hr +/− 3 hr ACI. *grnd^intr^*^a^ expression results in the rapid elimination of the clones from the disc within 48hr. Insets in **E** and **F** show the presence of the GFP-marked clones composed of 2-3 cells in each genotype at 24hr ACI. **G).** Expression of *Traf4^IR^* specifically in loser cells prevents their elimination from the wing disc. In contrast, *Traf4^IR^* expression in control clones does not alter their growth. **H).** Elimination of loser cells is also suppressed in the *Traf4^ex1^* null mutant background. **I).** Loss of *Traf6* by expression of *Traf6-RNAi* in the loser cells prevents their elimination. All clones grew from 48-96hr ± 2 hr AEL. Error bars show median and interquartile range. Statistical significance was determined by non-parametric Mann-Whitney tests. NS = not significant.

### Traf4 and Traf6 contribute to elimination of wild-type loser cells

In *Drosophila,* intracellular signaling via TNFRs is transduced through the TNF receptor-associated factors Traf4 (formerly known as dTraf1) and Traf6 (formerly dTraf2). TNF-TNFR association has been shown to activate both the JNK and the NF-κB signaling pathways, possibly because different intracellular signals can be transduced by different Traf adaptors (CHA *et al*. 2003). Traf4 physically associates with Wgn and Grnd, as well as Tak1, the Tak1-associated binding protein 2 (Tab2) and the Sterile-20 kinase, Misshapen (LIU *et al*. 1999; CHA *et al*. 2003; ANDERSEN *et al*. 2015). Expression of Traf4 is sufficient to induce JNK activity and cell death (CHA *et al*. 2003). Although signaling mediated by Traf4 is directed primarily through the JNK pathway, Traf4 has also been implicated in NF-κB-mediated responses (SHEN *et al*. 2001; AYYAR *et al*. 2007), and Traf4 is required to activate the Relish NF-κB factor for maintenance of proneural gene expression in wing discs (AYYAR *et al*. 2007).

We tested Traf4’s requirement in the elimination of WT loser cells by depleting its function in cell competition assays. Competition was suppressed when *UAS-Traf4^IR^*, encoding a *Traf4* inverted repeat (IGAKI *et al*.), was expressed specifically in the loser cells (Fig. 3G, Losers), whereas its expression did not affect the growth of control clones (Fig. 3G, Ctls). The elimination of loser cells was also suppressed in larvae carrying the severe *Traf4^ex1^* allele (CHA *et al*. 2003), and resulted in loser clones that were comparable in size to control clones in a non-competitive environment (Fig. 3H).

The Traf6 adaptor also associates with the intracellular domains of Wgn and Grnd (ANDERSEN *et al*. 2015). Unlike Traf4, ectopic expression of Traf6 is not sufficient to activate JNK signaling nor induce apoptosis, but it can participate in both JNK and NF-κB signaling (CHA *et al*. 2003; GEUKING *et al*. 2005). Traf6 functionally interacts with Pelle, the Toll pathway kinase, and in S2 cells Pelle and Traf6 cooperate in the activation of the NF-κB dorsal (SHEN *et al*. 2001). To examine the role of Traf6 in cell competition, we expressed the RNAi transgene *UAS-Traf6^IR^* (IGAKI *et al*.) in cell clones in both non-competitive and competitive contexts. Like loss of *Traf4*, knock-down of *Traf6* in the loser cells suppressed their elimination, allowing the clones to grow to the same size as non-competitive control clones (Fig. 3I). The loss of *Traf6*, like that of *Traf4*, had no obvious effect on cells in non-competitive control clones, indicating that its role was specific to the loser cells in the competitive context. Thus, the specific elimination of loser cells requires the function of both Traf4 and Traf6, consistent with their function downstream of Egr/Grnd within the loser cells.

### JNK signaling is activated, but has a minor role in elimination of WT loser cells

The MAP3K TGF-beta activated kinase Tak1 is critically required for transduction of JNK signaling downstream of Egr/Grnd activity (ANDERSEN *et al*. 2015). However, in previous studies, Tak1 was found to be dispensable in the elimination of loser cells in Myc– and in *Rp^-/+^*– mediated cell competition (MEYER *et al*. 2014). We confirmed these results here using two different *Tak1* null alleles that harbor point mutations in the kinase domain (VIDAL *et al*. 2001; DELANEY *et al*. 2006). We induced Myc cell competition in wing discs of male *Tak1^2^/Y* or in *Tak1^1^/Y* mutant larvae and found that neither mutation was sufficient to suppress elimination of the loser cells (Fig. 4A), confirming that loss of *Tak* does not interfere with the elimination of loser cells in this competitive context (MEYER *et al*.). This finding raises an interesting conundrum, since Tak1 is on the one hand required downstream of TNFR to activate JNK signaling (VIDAL *et al*.; DELANEY *et al*.; STRONACH *et al*.; ANDERSEN *et al*.), and on the other, it plays crucial roles in the activation of Rel in the immune response to pathogenic infection (SILVERMAN *et al*.; PARK *et al*.).

**Figure 4.**
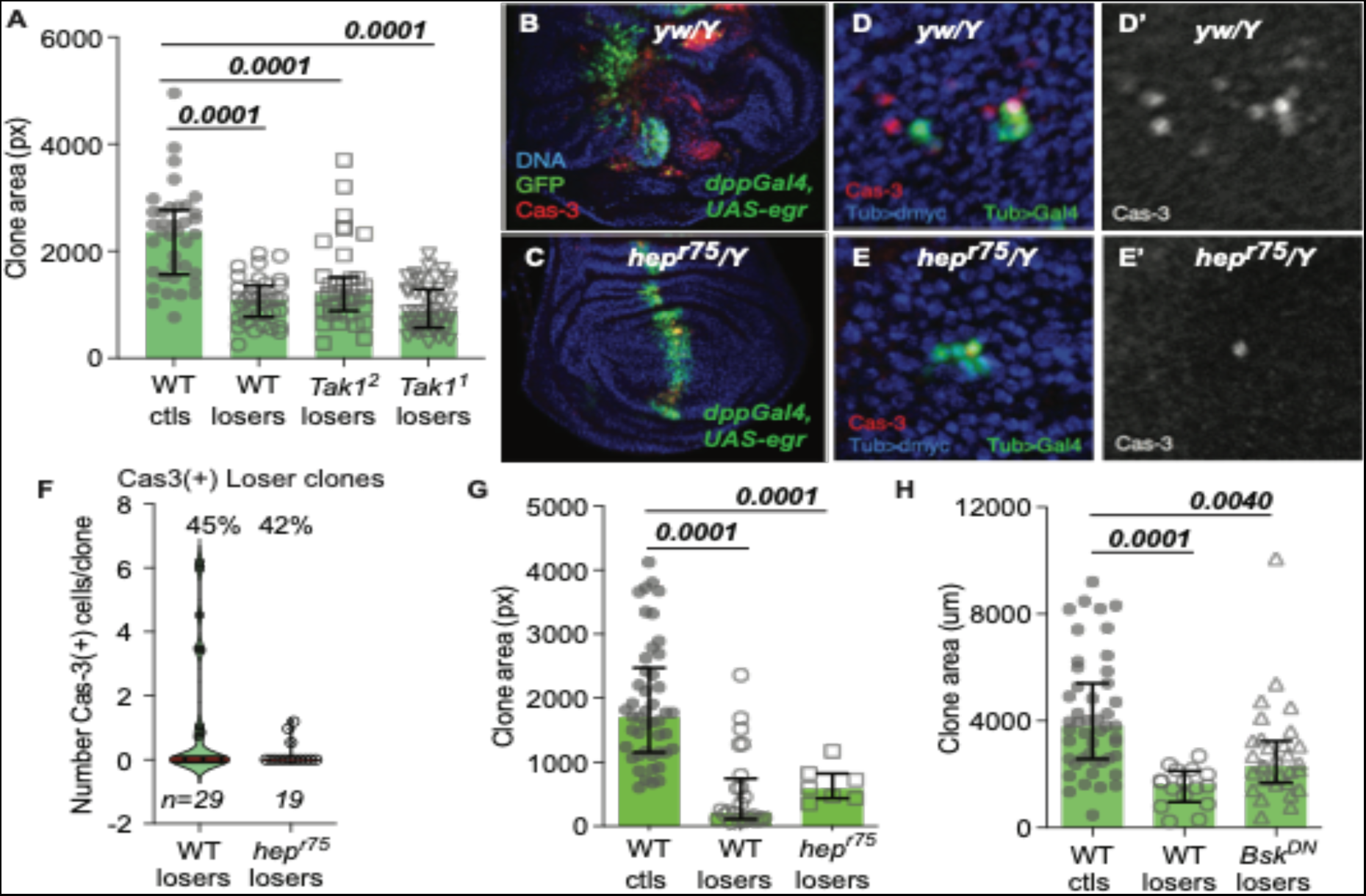
JNK signaling has a minor role in the elimination of WT loser cells. **A).** Loss of *Tak1* does not prevent cell competition. Two different null alleles of Tak1 were tested. Clones were scored in wing discs from male larvae (*Tak1/Y* and *WT/Y*). **B-C).** Expression of *UAS-egr* induces massive cell autonomous and non-cell autonomous cell death. *UAS-Egr* was co-expressed with *UAS-GFP* using *DppGal4* in WT **(B)** or *hep^r75^* wing discs **(C)**. Anti-activated Cas-3 (red) staining highlights dying cells. **C).** The *hep^r75^* allele blocks the majority of apoptosis induced by *UAS-egr.* Wing discs are from *WT/Y* males or *hep^r75^/Y* hemizygous male larvae. **D).** WT loser clones marked with GFP die at high frequency. Clones are 24hr ACI. (**D’).** Anti-activated Cas-3 staining. **E).** Cell death in loser wing disc clones is reduced in the *hep^r75^/Y* background compared to loser clones at 24hr ACI in *WT/Y*. **(E’).** Anti-activated Cas-3 staining. **F).** Violin plots with quantification of activated Cas-3 (+) cells/clone at 24hr ± 2hr ACI. The percent of loser clones containing Cas-3 (+) cells is shown at the top, and the number of Cas3 (+) cells per clone shown in the plots. n = number of clones scored per genotype (*WT/Y* and *hep^r75^/Y*). **G).** By 50 hr ACI, loser clones in a WT background are significantly smaller than controls, and *hep^r75^*does not prevent their competitive elimination. Clones were scored in wing discs from male larvae (*WT/Y* and *hep^r75^/Y*). **H).** Loser clones expressing *UAS-Bsk^DN^*are larger than WT loser clones, but are still significantly smaller than the controls. **I-J)**. Wing discs in both non-competitive (**I, I’**) and competitive (**J, J’**) backgrounds show JNK activity reported by TRE-dsRed. TRE activity is sporadic and independent of clones. Error bars in (**A), (G)** and **(H)** show median and interquartile range. Statistical significance was determined by non-parametric Mann-Whitney tests for each condition compared to controls; p values are relative to *yw* controls.

Ectopic expression of *egr* in a stripe bisecting the wing disc induces massive cell death, measured by immunostaining with an antibody against active Caspase-3 (Cas-3) (Fig. 4B). This death is mediated by Egr’s activation of the JNK pathway, as it is suppressed in larvae mutant for *hep^r75^*, which removes the function of the Hep/MKK7 protein (GLISE *et al*.) (Fig. 4B-C). Previous work reported that in cell competition assays in wing discs, only a fraction of loser cell death was suppressed in *hep^r75^*mutants (DE LA COVA *et al*. 2004), suggesting that JNK pathway activity is not sufficient to account for loser cell elimination. Indeed, in a wildtype background, 45% of wing disc loser clones contained Cas-3 positive cells at 24hr ACI (Fig. 4D, D’, F). Similarly, in *hep^r75^* mutant wing discs, 42% of loser clones contained Cas-3 positive cells, although on average, they contained fewer Cas-3 positive cells/clone (Fig. 4E, E’, F), suggesting loss of *hep* slightly reduced loser cell death, but did not wholly prevent it. Moreover, *hep^r75^* loser cells were ultimately eliminated from the wing discs as effectively as WT loser cells. By 50hr ACI, the median size of loser clones in *hep^r75^* mutant males was similar to that of loser clones in a WT background, and both were significantly smaller than non-competitive control clones (Fig. 4G). Consistent with these data, expression of a dominant negative form of the Jun kinase, Basket (Bsk), which functions downstream of Hep (RIESGO-ESCOVAR *et al*.; SLUSS *et al*.), in the loser clones allowed them to grow slightly larger, but they remained significantly smaller than controls (Fig. 4H). Altogether, our results indicate that although loss of *Tak1*, *hep* or *bsk* function can slightly diminish loser cell death, none of these essential intermediaries of the JNK pathway are sufficient to suppress the cells’ competitive loss from the wing disc.

### The function of Egr/Grnd in WT loser cells is independent of JNK activity

Together, our results establish that Egr activity, through its receptor Grnd, is required for the elimination of WT loser cells from the wing disc epithelium in Myc cell competition. However, our experiments also indicate that in the Egr/Grnd contribution to loser cell death, canonical JNK signaling downstream of the Traf4 and Traf6 adaptors is dispensable: loss of either *Tak1*, *hep* or *bsk* is insufficient to prevent loser cell elimination, despite their typical activation downstream of Egr/Grnd signaling and their critical requirement for JNK activity. These results suggest that in this competitive context, Egr/Grnd’s role in loser cell elimination is largely independent of JNK activity.

Previous work demonstrated that the NF-κB factor Rel is required for the expression of Hid and apoptosis of WT loser cells as part of the CCSM (MEYER *et al*. 2014; NAGATA *et al*. 2019; HOF-MICHEL *et al*. 2020). Here, in the same competitive context, our data indicate that the absence of either *egr* or *grnd* is also sufficient to prevent loss of loser cells. One way to explain these two findings is that in the loser cells, signaling from Egr/Grnd participates in or reinforces activation of Rel. If so, we reasoned that in this competitive context, activation of Egr/Grnd signaling in a *Tak1* mutant should not suppress loser cell death (i.e., the cells would still be eliminated). Conversely, in a non-competitive context, loss of *Tak1* would be predicted to completely block the canonical JNK signaling downstream of Egr/Grnd activation and allow the cells to survive (ANDERSEN *et al*. 2015). We tested these ideas by clonally expressing *grnd^intra^* in *Tak1^2^/Y* hemizygous male larvae, and quantifying the size of the clones in wing discs in both non-competitive control and competitive contexts. Female *Tak1^2^/+* larvae, which clonally expressed *grnd^intra^*but were heterozygous for *Tak1*, served as an internal control and exhibited massive death induced by the *grnd^intra^* expression that severely diminished larval growth (Fig. 5A) (ANDERSEN *et al*. 2015). By contrast, hemizygous *Tak1^2^/Y* males from the same cross were unable to transduce the *grnd^intra^*-mediated death signal, so the larvae grew normally (Fig. 5A). When cell competition was induced in WT male larvae, we observed the expected growth disadvantage in loser cells compared to non-competitive controls (Fig. 5B). In striking contrast, expression of *grnd^intra^* in either control or loser clones blocked their growth due to acute cell death (Fig. 3D-F and Fig. 5C, represented as ‘no clones’). As expected, the complete absence of *Tak1* completely prevented *grnd^intra^*-induced cell death in the non-competitive context, allowing the clones to grow to the same size as the WT controls (ctls, Fig. 5D vs. 5B). Moreover, in the competitive *tub>myc^STOP^>* context, lack of *Tak1* also allowed growth of *grnd^intra^* expressing clones (losers, Fig. 5D), however, these clones were significantly smaller than non-competitive control clones (losers, Fig. 5D vs 5B). This experiment demonstrates that although *Tak1* depletion blocked the massive death in the cells due to *grnd^intra^* expression, it did not prevent the cells from acquiring the “loser” fate that disadvantages them compared to control clones. This result confirms that Tak1 is not required in cell competition (MEYER *et al*.). Importantly, it also indicates that in the loser cells, the cell death induced through the CCSM still occurred, even while Tak1 depletion selectively prevented JNK-dependent cell death driven by *grnd^intra^*. Altogether, our results provide solid evidence that canonical JNK activity plays an insubstantial role in the death and elimination of WT loser cells (DE LA COVA *et al*. 2004).

**Figure 5.**
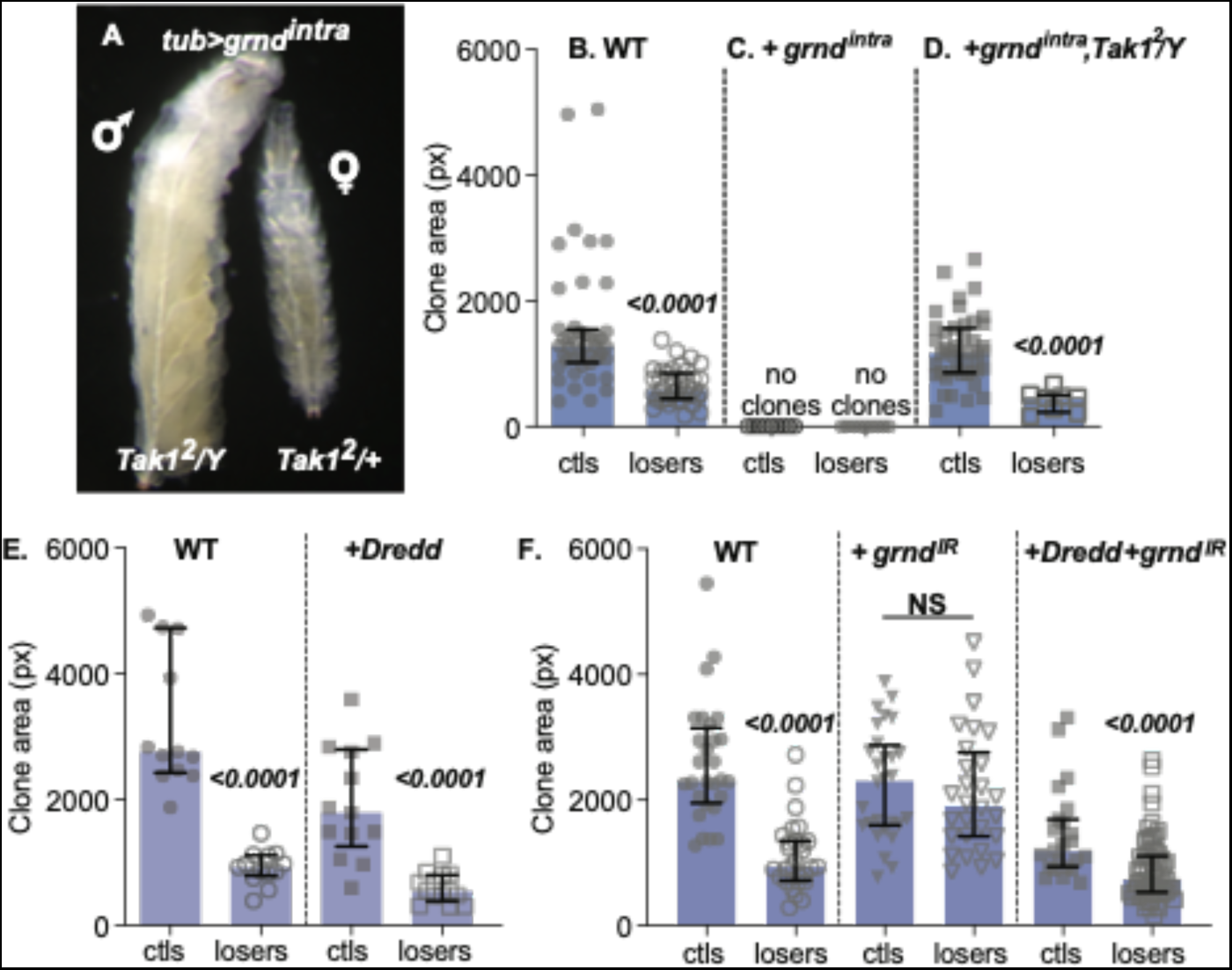
Evidence that in loser cells, Grnd/TNFR signaling diverges from JNK to the CCSM. **A).** Sibling *Tak1^2^* mutant male (left) and female (right) larvae clonally expressing cell *UAS-grnd^intra^* under *tubGal4* control. The complete loss of *Tak1* in hemizygous males abolishes the massive death induction due to expression of *Grnd^intra^*, while female *Tak1^2^/+* larvae remain susceptible to *Grnd^intra^ –*mediated cell killing and the consequent overall size reduction. **B-D).** *Tak1^2^* mutant prevents apoptosis induced by *grnd^intra^*in non-competitive control clones, but not cell competition-induced apoptosis in loser clones. **B).** Control and loser clones from *WT* males. **C).** Neither control nor loser clones expressing *grnd^intra^* grow, due to extensive cell death (see Fig. 3E**-F**); represented here as “no clones”. **D).** Under non-competitive conditions, *Tak1^2^/Y* prevents cell death induced by *grnd^intra^*expression in control clones (compare to **B**, *WT/Y* ctls), indicating that these cells die due to JNK-mediated signaling. In contrast, *Tak1^2^/Y* does not prevent the elimination of loser clones in cell competition (compare to **B**, *WT/Y* losers), indicating that JNK signaling is not required for their death. Clones grew for 48 ± 1.5 hrs. Error bars show median and interquartile range. Statistical significance was determined by non-parametric Mann-Whitney tests. **E).** Clones of WT loser cells (losers) clones are smaller than WT control clones (ctls) due to their competitive elimination. Right, Expression of *Dredd* in control clones induces cell death, reducing clone size (WT ctls vs +Dredd ctls). In the competitive context, *Dredd* expression in loser cells enhances their elimination (WT losers vs +Dredd losers). **F).** Knock-down of *grnd* via expression of *UAS-grnd-RNAi* (*grnd^IR^*) specifically in loser clones efficiently suppresses their elimination (+*grnd^IR^* losers), but does not alter the size of control clones (+*grnd^IR^* ctls). Co-expression of *Dredd* and *grnd^IR^* does not suppress cell death induced by *Dredd* expression in non-competitive controls (+*Dredd* +*grnd^IR^* ctls), nor does it suppress the elimination of loser cells (+*Dredd* +*grnd^IR^*losers). Clones were allowed to grow for 48 ± 1.5 hrs. Error bars show median and interquartile range. Statistical significance was determined by non-parametric Mann-Whitney (individual ctl/loser pairs) and Kruskal-Wallis (across genotypes) tests. NS = not significant.

### Dredd and Grnd have different genetic relationships in non-competitive versus competitive environments

How does Egr/Grnd function in the death of loser cells, if not via JNK activity? The function of the *Drosophila* caspase-8 ortholog Dredd is required upstream of Rel for loser cell death (MEYER *et al*. 2014). Dredd is also required for immune signaling by the IMD pathway, and over-expression of Dredd is sufficient to induce proteolytic activation of Rel and leads to Rel-dependent gene expression in immune competent tissues (STOVEN *et al*. 2003; ZHOU *et al*. 2005). In wing discs, *Dredd* expression leads to Rel-dependent cell death (MEYER *et al*. 2014). Moreover, the death of WT loser cells in the competitive context is dominantly suppressed in *Dredd/+ Rel/+* double heterozygous mutants, suggesting that the loser cells have increased sensitivity to Dredd activity (MEYER *et al*. 2014). Notably, however, Dredd is not required for cell death induced by canonical JNK signaling downstream of Egr (IGAKI *et al*. 2002; MORENO *et al*. 2002b).

To further explore the relationship between Egr/Grnd/JNK signaling and the CCSM, we tested for genetic interactions between JNK signaling and Dredd. Expression of *UAS-Dredd* in cell clones in a non-competitive context leads to Rel-dependent apoptosis (MEYER *et al*.) and significantly reduces clone size compared to non-expressing controls (Fig. 5E, ctls). We tested whether JNK signaling plays a role in this process by depleting Grnd from the cells. In the competitive *tub>myc^STOP^>* context, where Dredd is already active in the loser cells, expression of *UAS-Dredd* enhances their elimination (Fig. 5E, losers) (MEYER *et al*.). Furthermore, expression of *grnd* RNAi (*grnd^IR^*), like the complete loss of *grnd* (Fig. 2E), fully suppresses the competitive elimination of loser cells without altering the growth of control clones (Fig. 5F, +*grnd^IR^*).

Since loss of *grnd* prevents Egr-induced activation of JNK signaling (ANDERSEN *et al*. 2015), we tested whether Dredd-induced cell killing required input from Egr/Grnd activity. We postulated that by preventing Egr-induced signaling via *grnd^IR^*, cell death induced by *UAS-Dredd* expression would be detached from input by JNK activity. Indeed, in the non-competitive context, *grnd* knock-down was unable to suppress the clone size reduction caused by *Dredd* expression (Fig. 5F, *Dredd* +*grnd^IR^*ctls). This indicates that in a non-competitive context, Dredd-induced cell death does not require input from Grnd. In contrast, in the *tub>myc^STOP^>* competitive context, loser cells that expressed *Dredd* + *grnd^IR^* retained the loser fate, as the cells were still eliminated and the clones significantly smaller than their cognate controls (Fig. 5F, *Dredd* +*grnd^IR^* losers). These results provide strong evidence that in both the non-competitive and competitive contexts, the activity of Dredd does not require JNK signaling to induce cell death.

Together, these experiments argue that in the WT loser cells, Egr/Grnd activity reinforces the CCSM-mediated cell death response that restricts growth of the loser clones. However, they also argue that Egr/Grnd can participates in two parallel signaling pathways within the loser cells: one that is Tak1 independent and reinforces the CCSM, in which cell death is mediated by Dredd and Rel (MEYER *et al*.); and another that directs signaling through the canonical JNK pathway mediators and activates JNK reporters (DE LA COVA *et al*.; MORENO AND BASLER), but provides only a minor contribution to loser cell death. We speculate that the enhancement of loser cell elimination conveyed by UAS-*Dredd* expression (Fig. 5E, *+Dredd*), which appears slightly diminished in the absence of *grnd* (Fig. 5F,*+Dredd+grnd^IR^* losers), could be interpreted as evidence of parallel, additive signaling. Our results imply that Egr/Grnd-induced JNK signaling and Egr/Grnd-assisted CCSM signaling function in parallel during cell competition, but that JNK signaling plays a minor role in the loser cells. Moreover, our results make it clear that complex signaling inputs are deployed in the loser cells. This work thus helps to illuminate the qualitatively different survival and death cues that WT cells growing in the *tub>myc^STOP^>* context (ie, loser cells) are subject to compared with genotypically-identical WT cells in a non-competitive environment.

## Discussion

Although the JNK pathway is activated during Myc cell competition, the relative contribution of this pathway to the elimination of the WT loser cells, and how JNK signaling and the CCSM interact, have remained unclear. We have attempted to understand these interactions with a series of genetic experiments. We establish that loss of Egr/TNF or Grnd/TNFR suppresses the elimination of loser cells in the competitive *tub>myc^STOP^>* background, indicating that each factor is functionally required in the process. Although Egr is known to kill cells via JNK signaling, our experiments indicate that JNK pathway activity has only a minor role in the death of WT loser cells (Fig. 4A-G) (DE LA COVA *et al*. 2004). Our results argue that in Myc cell competition assays, signaling in the loser cells by Egr/Grnd bifurcates upstream of Tak1, to facilitate and/or potentiate the activity of the CCSM in a process that is independent of downstream JNK pathway activity. Collectively, our experiments bolster earlier findings that JNK signaling is not sufficient to cause the elimination of loser cells in this competitive context (DE LA COVA *et al*. 2004), in contrast to the NF-kB Rel, which is critical for loser cell death (MEYER *et al*. 2014; NAGATA *et al*. 2019).

Grnd is highly and ubiquitously expressed in wing discs, whereas the TNFR Wgn expression appears to be more restricted. The expression of *wgn* in the WP (Fig. 2A), where cells are most severely subject to Myc super-competition (ALPAR *et al*.), raises the possibility that Wgn and Grnd could act semi-redundantly in that region to promote loser cell elimination. However, the loss of Wgn is inconsequential in loser cells (Fig. 2C-D) whereas loss of Grnd prevents their competitive elimination (Fig. 2E). Our data therefore indicate that Grnd functions as the primary Egr receptor in wing discs in this experimental context. Egr itself is expressed in multiple tissues in the larva, and can be produced as a membrane bound protein or in a soluble form after proteolytic processing (KAUPPILA *et al*. 2003; NARASIMAMURTHY *et al*. 2009). Although Egr is highly expressed in the AMP cells that reside under the notum region of the wing disc (Fig. 1C), it is undetectable in most epithelial cells of the disc. We were also unable to detect any *egr* mRNA expression in either loser (or winner) cells in the competitive context (Fig. 1I-I’), suggesting that *egr* expression is not an autocrine product of CCSM activation in the loser cells. Instead, we speculate that Egr, which appears to be constitutively present in the circulating hemolymph (AGRAWAL *et al*.), is produced and secreted by another tissue into circulation, from which it then associates with Grnd in the disc epithelial cells (Sharma Singh et al, in prep). It is possible that circulating Egr signals in discs via Grnd to induce the low, tonic JNK activity observed in the wing disc. If so, perhaps it serves to “prime” the loser cells to the effects of the CCSM.

The possibility that Egr is secreted remotely has precedence in previous work, in which Egr, synthesized in the larval fat body (FB), can lead to signaling in cells of the brain or wing disc (AGRAWAL *et al*.; DE VREEDE *et al*. 2022). Grnd is present on the apical membranes of wing disc cells, which face inwards toward the disc lumen, and is also found in intracellular vesicles {Palmerini, 2021 #172}. Recent work suggested the apical localization of Grnd prevents easy access to secreted Egr unless cell polarity is lost, as in *scribbled* mutant discs, in which case Grnd mislocalizes to the basolateral membrane (DE VREEDE *et al*. 2022). Such mislocalization does not occur in either the loser cells or in the Myc expressing super-competitor cells in the context of the competitive assay (see Fig. 2B), suggesting there is a different mechanism for the association of Egr-Grnd in this context.

Regardless of its tissue source, our results suggest a mechanism for how Egr contributes to the death of loser cells in wing discs. We postulate that the binding of Egr to Grnd on wing disc cells contributes to activation of a low tonic level of JNK activity in wing discs (Fig. 1F-G). We speculate that once Egr binds to and activates Grnd on wing disc cells, this information is transmitted within the cells in two ways. First, our results suggest that in loser cells, Egr/Grnd activity potentiates CCSM activity – possibly via Dredd –-to reinforce their death (MEYER *et al*.). At the same time, Egr/Grnd also activates the canonical JNK pathway, which may provide a minor contribution to their demise (DE LA COVA *et al*.). How do winner cells escape the cell death induced by the CCSM and JNK activity? We suggest two non-exclusive possibilities. First, given the propensity of increased Myc expression to suppress Egr-JNK signaling (HUANG *et al*. 2017), cells harboring the *>myc^STOP^>* cassette may be less susceptible to the death inducing role of JNK activity. Second, although the *>myc^STOP^>* expressing winner cells provide the cues that trigger CCSM activation, the CCSM’s killing activity is restricted to the loser cells (MEYER *et al*.; ALPAR *et al*.), thus biasing any augmentation of its activity to the WT cells.

Although the exact mechanism by which the Egr/Grnd signal contributes to the CCSM remains to be determined, our data suggest that it bifurcates downstream of the Traf adaptors, above Tak1. Both Traf4 and Traf6 are required in our assays for loss of the loser cells, whereas the absence of Tak1 does not affect the outcome of cell competition, even while it abrogates JNK signaling. These findings provide another example of productive cross-talk between TNF/TNFR and NF-kB signaling pathways, which occurs in both the mammalian and *Drosophila* innate immune responses and also diverges at the level of Tak1/Tab2 (MYLLYMAKI AND RAMET 2013). Our genetic interaction tests with Grnd and Dredd suggest that downstream of Egr/Grnd activation, Traf4/6 could facilitate and/or enhance Dredd activity in the loser cells, thereby augmenting Rel-dependent, loser-specific gene expression and cell death. Intriguingly, although Dredd and Rel function in the CCSM, neither Tak1 nor the IKK complex are required for loser elimination in Myc cell competition (MEYER *et al*.), perhaps illustrating how these pathways can be repurposed for different biological tasks.

What is the nature of the interactions between the activities of the JNK pathway and the CCSM? In the immune response, JNK activation is short-lived (SLUSS *et al*. 1996), in part because of feedback from Rel that dampens JNK signaling by promoting the degradation of Tak1 (PARK *et al*. 2004; ZHOU *et al*. 2005). Alternatively, Traf4 is known to activate the dorsal/NF-κB pathway (LIU *et al*. 1999; SHEN *et al*. 2001), thus Egr/Grnd signaling could lead to dl/Dif activity as well as Rel, strengthening the outcome in the loser cells, and even potentially taking the form of a feed-forward activity that stabilizes JNK activity in the loser cells. Interestingly, the elimination of loser cells appears to be stochastic, with some loser cells falling to the death-inducing signals more easily than others. The probability of loser cell death are likely influenced by several independent factors, such as accumulation of the proapoptotic factor Hid, which in turn would be influenced by inputs to the *hid* locus via the activities of the CCSM and JNK signaling; tonic JNK activity in the loser cells could even prime them for death by the CCSM.

We note that the intersection of the CCSM and JNK signaling that we describe here may be unique to the context of Myc-induced competition (and possibly competition between WT and *Rp^-/+^* cells, where both pathways have also been documented (MORENO *et al*.; TYLER *et al*.; MEYER *et al*.)). Competition between the polarity deficient *scrib* mutant cells and WT cells also utilizes Egr-JNK signaling to eliminate *scrib* loser cells, but this is Tak1-dependent, and as mentioned, also requires Wgn (IGAKI *et al*. 2006). Future studies will be important to sort out these differences. Nonetheless, our results help to explain the basis of the relative contributions of the CCSM and JNK activity to Myc-induced cell competition. In addition, they add to the growing body of work that illustrates the remarkable signaling flexibility that occurs between cells in competitive contexts, allowing cells to tune their response to a given cellular or environmental context.

## ACKNOWLEDGEMENTS

We thank P. Leopold, B. Lemaitre, M. Miura, M. Milan, T. Igaki, and D. Bohmann for *Drosophila* strains, P. Leopold for antibodies. We are grateful to members of the Johnston lab for helpful advice, C. Cary, P. Guevarra, D. Miranda and J. Park for excellent technical support. We are indebted to the BDSC (supported by NIH P40OD018537), DGRC (supported by NIH 2P40OD010949) and FlyBase (ATTRILL *et al*.) for their valuable resources. Funding for this work was provided by grants from the NIH (R01GM078464 and R35GM131871) and the NCI (R01CA192838) to L.A.J.

## AUTHOR CONTRIBUTIONS

All authors conceived and designed experiments. A.K., A.S.S., C. de la C. and M. Z. conducted experiments and analyzed results. A.K., A.S.S., C. de la C. and L.A.J. contributed to writing the paper. L.A.J. supervised the project and provided grant support.

## FIGURE LEGENDS

(In all graphs, clone area is measured in either pixels or microns, as indicated. Clones labeled control (ctl) were produced by FLP-out of the *act>CD2^STOP^>Gal4* cassette and clones labeled losers were produced by FLP-out of the *tub>myc^STOP^>Gal4* cassette).

**Supplementary Figure 1.**
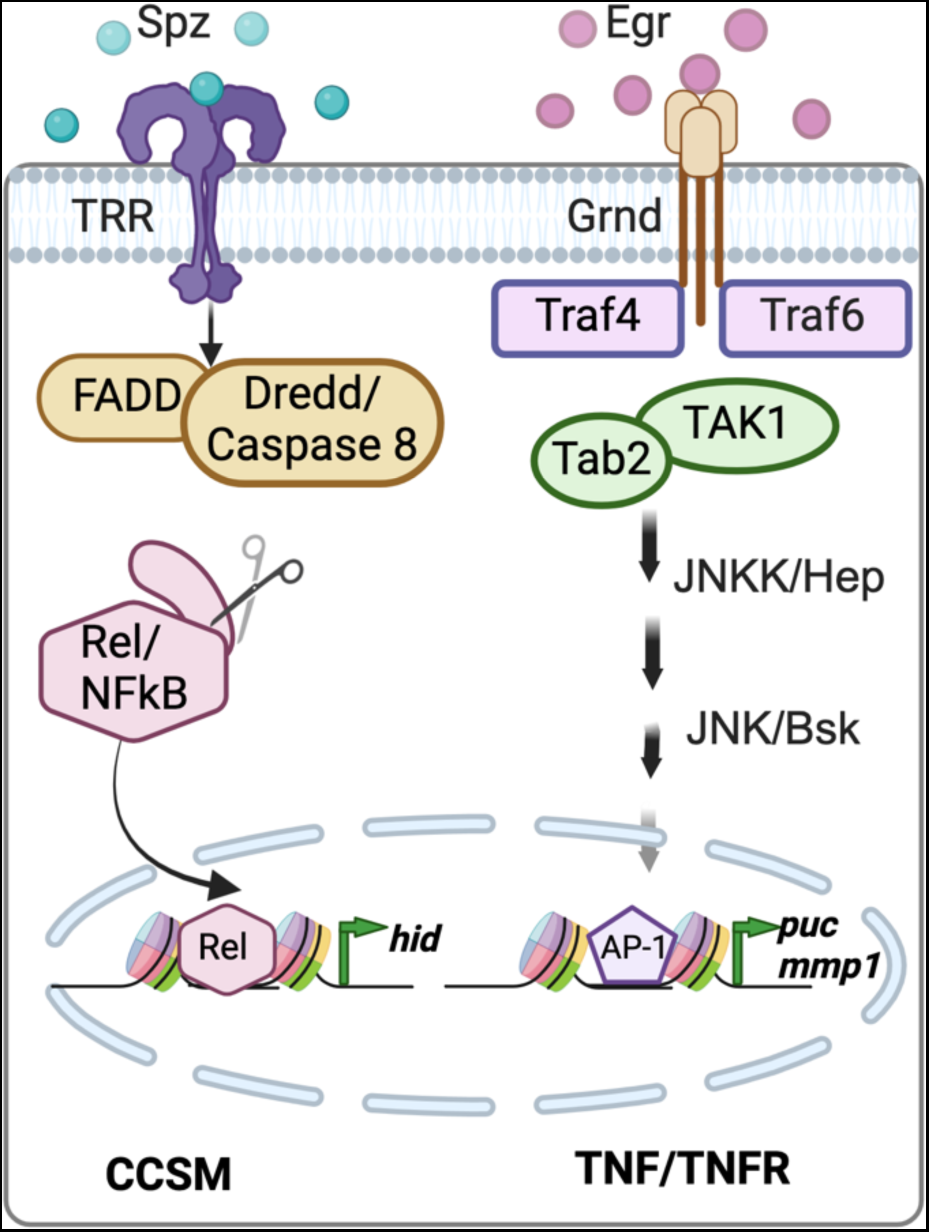
Schematic models of the CCSM and TNF/TNFR signaling pathways. **Left,** model of the CCSM pathway. The model (based on data from (MEYER *et al*. 2014) and (ALPAR *et al*.)) proposes that the Spz ligand, which is activated via processing by SPE (MEYER *et al*. 2014) associates with Toll 1 (Tl), possibly as a heterodimer with Toll-8 (Tollo). Toll-2 (18w), Toll-3 (Mst-prox) and Toll-9 are also genetically required for the elimination of loser cells in Myc cell competition, but their roles remain to be determined (MEYER *et al*. 2014; ALPAR *et al*. 2018). Transduction of the Spz/Tl association requires the function of the TIR domain protein Ect4 (not shown), FADD, Caspase-8/Dredd, and Rel to lead to loser cell death, and Rel requires cleavage by Dredd to induce apoptosis of the WT loser cell (MEYER *et al*. 2014). The direct targets of Rel in cell competition remain to be determined, but the *hid* gene is activated in a Rel-dependent manner. The anti-microbial peptide genes that Rel activates in innate immunity are not induced during cell competition (MEYER *et al*. 2014). **Right,** model of conventional TNF/TNFR signaling in *Drosophila* (COLOMBANI AND ANDERSEN 2023). The TNF Egr associates with Grnd/TNFR on the plasma membrane of the cell. Traf4 and/or Traf6 function as adaptors within the cell, and activate the core kinase cascade of the JNK pathway that included Tak1/MAPKKK, Tab2, Hep/JNKK, and Bsk/JNK. Targets of the JNK pathway include the genes encoding the metalloprotease Mmp1 and the phosphatase Puckered, among others.

